# Gut microbiome connectivity drives mass host colonization and buffers against antibiotic-induced collapse

**DOI:** 10.64898/2026.03.05.709772

**Authors:** Maria S Bañuelos, Bryan K Lynn, Mirjam Zünd, Rose Faucher, Calvin Trinh, Maria Rebolleda-Gómez, Travis J Wiles

## Abstract

Microbial dispersal continually shapes gut microbiomes, seeding communities during assembly and replenishing them after perturbation. Dispersal’s effect depends on its strength: low dispersal introduces stochasticity and drives divergence, whereas high dispersal increases mixing and promotes convergence. Although this pattern manifests across host–microbe systems, the mechanism by which dispersal alters colonization dynamics to generate such contrasting outcomes remains unclear. Identifying factors that toggle these dualistic effects is essential for predicting and controlling gut microbiome assembly and stability. We addressed this problem in gnotobiotic larval zebrafish, which is a tractable vertebrate model that enables experimental control and quantification of gut bacterial dispersal. To dissect the roles of dispersal, we defined the relationship between inoculation dose and colonization frequency for a model *Vibrio cholerae* isolate under different dispersal regimes. Strikingly, a dose yielding only 50% colonization in isolated hosts—the CD50—produced nearly universal colonization during co-housing, despite identical host and bacterial densities. Measurements of intestinal growth, carrying capacity, shedding, and environmental persistence were used to construct a quantitative colonization–dispersal model that explained the basis for this shift in colonization outcomes. At the CD50, colonization is inherently probabilistic, but once the first hosts become colonized, they reseed the environment and amplify secondary exposures, creating a feedback loop that rapidly transforms individual-level stochasticity into widespread colonization. We further show that this feedback operates in complex microbiomes, where larger host groups—with more opportunities for recolonization—buffer communities against antibiotic-induced collapse. Together, our findings demonstrate that dispersal regimes are not fixed but dynamically shift as hosts become increasingly connected. By revealing how dispersal and interhost transmission drive mass colonization and stabilize gut communities, our work identifies microbiome connectivity as a central mechanism governing gut microbial assembly and resilience.

## Introduction

Human and animal gut microbiomes are among the densest and most complex microbial ecosystems on Earth. These resident communities perform vital functions for their hosts that include cueing developmental pathways, modulating immunity, regulating metabolism, defending against pathogens, and preventing chronic diseases [1–6]. When microbiome assembly is disrupted or the community is damaged, these critical services can be compromised. Consequently, understanding how to preserve microbiome integrity and function is a major focus in the field. However, microbiome assembly and stability depend on numerous factors that interact dynamically, in some cases amplifying or overriding each other [7–12]. Identifying which factors can serve as effective therapeutic targets remains a major challenge.

A prevailing view of microbiome assembly is that it is determined by nonneutral, niche-based processes rooted in host biology and co-evolved host–microbe relationships [7,13,14]. Host traits such as gut physiology, immune activity, and mucosal architecture create a complex landscape of chemistries and microhabitats that select for or filter compatible bacterial taxa [15–20]. Microbial traits common to pathogens, pathobionts, and commensals—including motility, adhesion, immune evasion, and metabolic activity—further modulate colonization success [21–24]. Together, host and microbial factors interact to structure assembling consortia, but they do not act in isolation.

Dispersal controls the order and rate with which microbial species arrive in the gut, strongly influencing priority effects, in which early colonizers competitively exclude invaders [25,26]. In this way, low to moderate levels of dispersal introduce stochasticity, interacting with niche-based filtering to produce divergent assembly trajectories and the pronounced variation seen in microbiome composition across individuals [25,27,28]. Yet high levels of dispersal promote microbiome homogenization, as seen in hosts that live in geographic proximity or closely connected social groups, indicating that microbial exchange in certain contexts can overwhelm local filtering [10,29,30]. Landmark studies in mice revealed that shared environments enable the transfer of gut consortia that can induce pathological host phenotypes, including intestinal inflammation and metabolic syndromes [31–33]. In zebrafish—a vertebrate model ideally suited for testing predictions about interhost dispersal—co-housing wild-type and immune-deficient animals homogenizes microbiomes and enriches for taxa encoding traits that facilitate spread, such as motility and chemotaxis [10,34]. These findings challenge conventional views of gut microbiomes as closed ecological systems by showing that dispersal can exert a dominant influence on community membership. However, the mechanisms by which dispersal shapes colonization dynamics and toggles communities between divergent, individualized states and convergent, homogenized assemblages remain poorly understood.

In the present study, we set out to clarify the mechanistic roles of dispersal, and more specifically interhost transmission, during gut microbial community assembly. Recent work in invertebrate models has shown that gut bacterial colonization is inherently stochastic and dose-dependent such that exposure to intermediate doses yields only chance colonization across hosts even under identical conditions [35–37]. Host and microbial traits can shift colonization–dose relationships, but we hypothesized that dispersal could also further modify the distribution of colonization probabilities among hosts within a group.

We tested this hypothesis in larval zebrafish, where we found that a bacterial dose capable of yielding only a 50/50 colonization frequency in isolated hosts surprisingly achieves a near 100% colonization rate during co-housing. A quantitative colonization–dispersal model revealed that once a single host becomes colonized, it seeds the environment, increasing exposure probabilities for nearby hosts. This dynamic essentially transforms an initially probabilistic process into a near certainty at the group level. We further demonstrate that this phenomenon extends beyond initial assembly, revealing that animals belonging to larger group sizes—which contain a larger number of reservoirs for recolonization—display increased microbiome stability during antibiotic treatment. Together, our results uncover a mechanistic basis for how dispersal overcomes local filtering and bottlenecks to drive mass colonization, highlighting interhost transmission as a key lever for managing and engineering microbiomes.

## Results

### Interhost transmission sets the probabilistic colonization threshold

To investigate how dispersal and interhost transmission shape colonization frequencies, we adapted an established gnotobiotic larval zebrafish model to measure dose-dependent colonization of hosts by a single bacterial strain. For these studies, we selected *Vibrio cholerae* ZWU0020 (hereafter “*Vibrio*”), which is a well-characterized strain that represents lineages prevalent in zebrafish gut microbiomes in both laboratory and natural settings [21,38–40].

Dispersal is difficult to experimentally model because it requires controlling not only the movement of microbes among hosts, but also their growth and persistence in the surrounding environment. A critical feature of our experimental design was therefore to ensure that colonization outcomes reflected initial encounter probabilities, rather than subsequent environmental overgrowth. To achieve this, fish were transferred to new sterile embryo media prior to inoculation of *Vibrio* into the water column. In conditioned water previously inhabited by fish, *Vibrio* proliferates rapidly, saturating the environment and obscuring dose-dependent colonization responses (S1A–C Fig). In contrast, in unconditioned sterile media, *Vibrio* viability quickly decays (S1D Fig), enabling us to better control initial environmental abundances and relate them to colonization outcomes.

Using this setup, isolated animals display a clear dose-dependent colonization pattern at 1 day post-inoculation (dpi) (Fig 1A). Colonization never occurred at low inocula (10^2^ bacteria/mL) and was universal at high doses (10^7^ bacteria/mL). The dose at which approximately half of the animals became colonized (∼10^5^ bacteria/mL) defines the colonization dose 50 (CD50) and therefore the probabilistic threshold for colonization success. Notably, fish colonized under this regime had a median gut abundance of ∼10^4^ bacteria/gut, roughly tenfold lower than in conditioned water (S1A Fig), suggesting that water quality may alter *Vibrio* colonization traits or that environmental overgrowth leads to a persistent, high dispersal pressure capable of overshooting carrying capacity.

**Figure 1.**
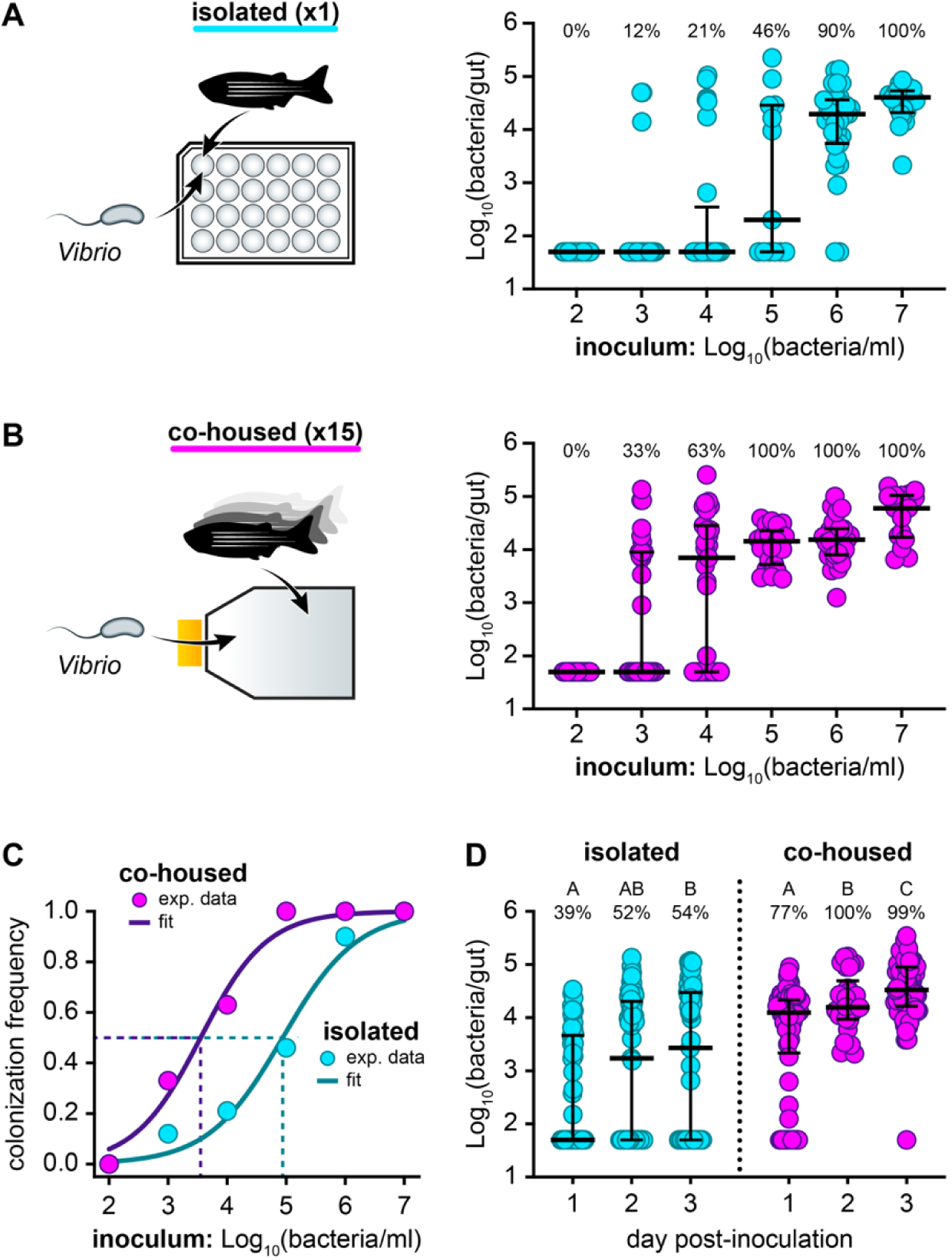
Interhost transmission sets the probabilistic colonization threshold. (A) Left: Isolated larval zebrafish were colonized with *Vibrio* in individual wells of a multiwell plate at a density of 1 fish/mL. Right: *Vibrio* gut abundances in isolated animals 1 day post-inoculation (dpi) with a range of bacterial doses. Sample sizes (left to right): 21, 25, 29, 13, 31, 21. (B) Left: co-housed larval zebrafish were colonized with *Vibrio* in tissue culture flasks at a density of 1 fish/mL and 15 animals/flask. Right: *Vibrio* gut abundances in co-housed animals at 1 dpi across the dose range used in A. Sample sizes (left to right): 29, 39, 27, 23, 30, 17. Percentages in both A and B indicate the fraction of colonized animals. (C) Estimation of CD50 for isolated and co-housed animals. Points indicate colonization frequencies from panels A and B; curves show best-fit Hill functions describing the dose-response relationship. Dashed lines mark CD50 values: isolated CD50 = 8.7 × 10^4^ bacteria/mL, co-housed CD50 = 3.6 × 10^3^ bacteria/mL. (D) *Vibrio* gut abundances in isolated and co-housed animals inoculated at the CD50 for isolated animals (10^5^ bacteria/mL). Guts were sampled at 1–3 dpi. Bars denote medians and interquartile ranges. Letters signify statistical groupings determined a Kruskal-Wallis and Dunn’s multiple comparisons test (isolated: *p*=0.0219; co-housed: *p*<0.0001). Sample sizes for isolated fish (left to right): 58, 61, 57. Sample sizes for co-housed fish (left to right): 80, 30, 100. Percentages indicate the fraction of colonized animals. In all panels, animals are considered colonized when they contain ≥ 10^3^ bacteria/gut.

To test whether co-housing alters colonization probabilities, we colonized fish housed in groups of 15 while maintaining the same density (1 fish/mL) and inoculating with the same dose range used for isolated animals (10^2^–10^7^ bacteria/mL). Under these conditions, co-housed fish showed markedly higher colonization rates, reducing *Vibrio*’s apparent CD50 by over an order of magnitude compared to isolated animals (Fig 1B). Based on these data, we estimated CD50 values of 8.7×10^4^ bacteria/mL for isolated animals and 3.6×10^3^ bacteria/mL for co-housed animals (Fig 1C). We further found that this shift is stable over time, with colonization frequencies and gut abundances for both groups increasing only marginally beyond 1 dpi, indicating that host colonization is determined rapidly following inoculation (Fig 1D).

### Host bacterial shedding promotes mass colonization of co-housed animals

Because environmental bacterial abundance strongly influences zebrafish colonization, we next asked whether bacterial levels in the water could reveal mechanisms underlying colonization dynamics during co-housing. To address this, we quantified bacterial abundances in the water column of isolated and co-housed animals across a range of inoculum doses. At 1 dpi, median water abundances scaled with inoculum dose in both housing conditions, but striking differences emerged in relation to colonization outcomes. For isolated animals that remained uncolonized, water bacterial abundances collapsed to near the detection limit, whereas colonized hosts were associated with markedly higher levels (Fig 2A, left). Co-housing mirrored the trend of colonized isolated animals, exhibiting robust water bacterial abundances that closely tracked inoculum dose and host colonization status (Fig 2A, right).

**Figure 2.**
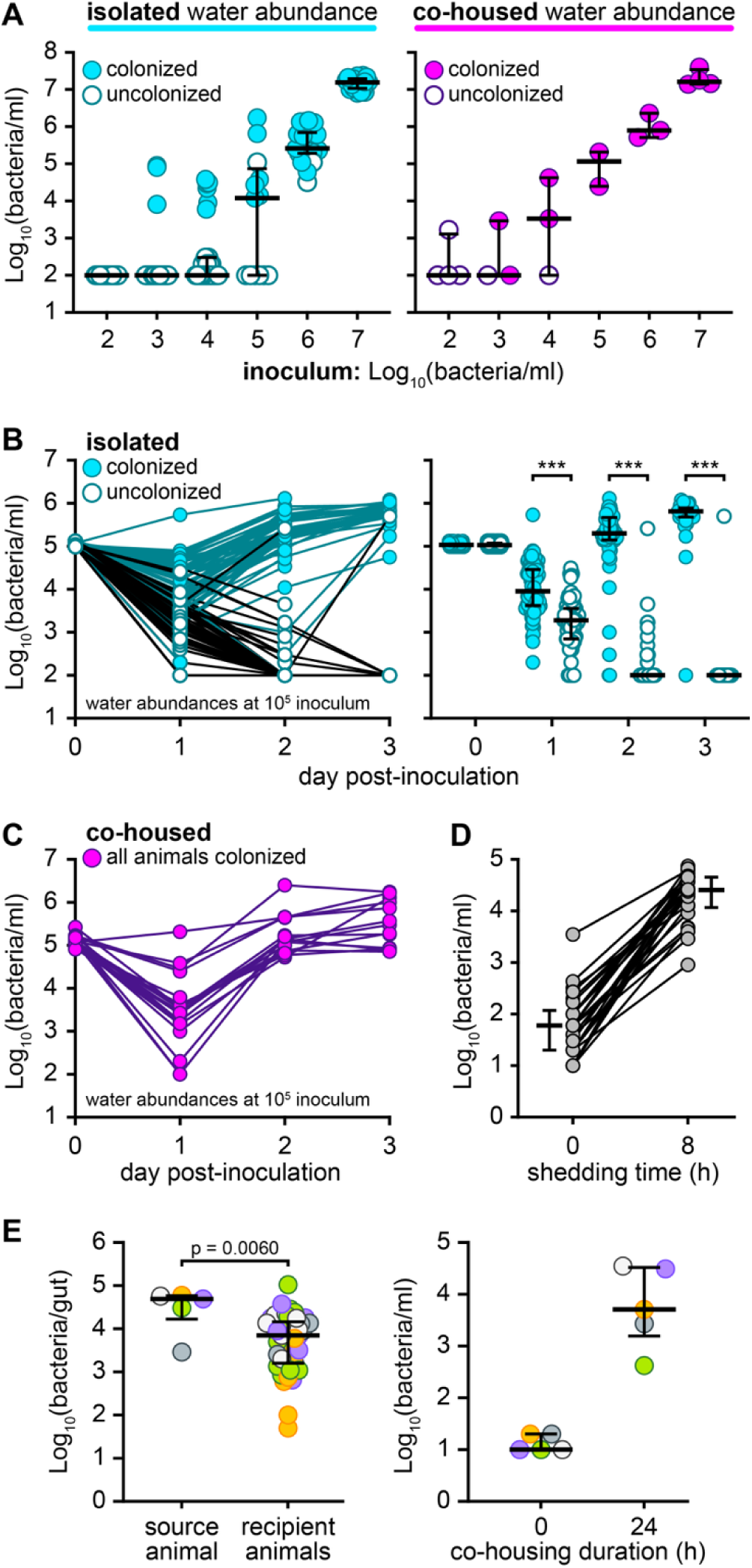
Host bacterial shedding promotes mass colonization of co-housed animals. **(A)** *Vibrio* water abundance from isolated (left) and co-housed (right) fish at 1 dpi, derived from the same experiments shown in Fig 1A and B. Cyan circles represent water associated with colonized hosts; empty circles represent water from uncolonized hosts. Bars indicate medians and interquartile ranges. Sample sizes for water abundances from isolated animals (left to right): 21, 26, 29, 13, 31, 22. Sample sizes for water abundances from flasks of co-housed animals (left to right): 4, 3, 3, 2, 3, 4. (B) Left: Water abundance trajectories from 0–3 dpi in isolated animals inoculated at the CD50 (data derived from experiment shown in Fig 1D). Each line represents an individual fish. Cyan circles denote water samples from colonized animals; empty circles denote samples from uncolonized animals. Right: Replotted trajectories from isolated animals to compare abundances between colonized and uncolonized animals at each time point. Sample sizes (left to right): 89, 87, 89, 87, 63, 55, 31, 26. Statistical significance was assessed by Mann–Whitney test; *p* < 0.001. (C) Water abundance trajectories from 0–3 dpi for co-housed animals inoculated at CD50 for isolated animals (data derived from experiment shown in Fig 1D). Each line represents water from a single flask of co-housed animals; all flasks contained colonized individuals. Sample size = 24. (D) Host bacterial shedding over an 8h interval. Lines connect paired measurements from the same fish. Adjacent bars indicate medians and interquartile ranges. Sample size = 26. Median shedding rate: 3.16 × 10^3^ bacteria/h ± 2.60 × 10^3^. (E) Gut (left) and water (right) abundances following the introduction of colonized source fish into groups of germ-free recipient animals. Bars indicate medians and interquartile ranges. Points of the same color represent samples originating from the same source–recipient group. Sample sizes for source and recipient fish: 5 and 51, respectively. Sample sizes for water abundances = 5 each.

To further resolve colonization dynamics, we plotted water bacterial abundances and host colonization status over time using data derived from the experiment in Fig 1D, which was performed with the approximate CD50 for isolated animals (10^5^ bacteria/mL). By 1 dpi, water abundances for isolated animals were already predictive of colonization outcome and became increasingly definitive at 2 and 3 dpi (Fig 2B). Both colonized and uncolonized animals exhibited an initial steep decline in water bacterial abundance, but the decline was less pronounced for colonized animals, and levels rebounded sharply by 2–3 dpi. In co-housed animals, water bacterial abundances also dropped initially but consistently recovered to high levels over time (Fig 2C). Importantly, control experiments confirmed that water conditioning by fish did not account for *Vibrio* recovery at later time points (S1E Fig).

Together, these data begin to build a model in which early declines in *Vibrio* water abundances create a narrow window for colonization. If no host is colonized, environmental populations continue to decay, making colonization impossible. In contrast, successful colonization transforms hosts into bacterial reservoirs that shed cells back into the water, replenishing environmental abundances. In this way, a colonization–dispersal feedback loop is formed that stabilizes bacterial populations and enables colonization of naïve co-housed hosts.

To evaluate whether host shedding occurs at a high enough rate to provide environmental rescue consistent with our model, we quantified *Vibrio* shedding from individual colonized fish over an 8-hour period (Fig 2D). Water bacterial abundances increased over time, corresponding to a median shedding rate of approximately 3.16 x 10^3^ cells per hour (SD ± 2.60 x 10^3^), a value consistent with intestinal clearance dynamics reported for larval zebrafish [21,41]. To directly assess whether shedding promotes colonization of naïve hosts, we co-housed a single colonized animal with 14 germ-free recipients. After 24 hours, nearly all recipients were robustly colonized, with gut bacterial loads approaching those of the source host concurrently with high water abundances (Fig 2E). These results demonstrate that even a single colonized host can act as a potent source of bacteria driving mass colonization.

### A quantitative colonization–dispersal feedback model describes rapid mass host colonization

Building on our experimental findings, we constructed a quantitative model (Fig 3A) grounded in empirical data to validate whether a colonization–dispersal feedback loop during co-housing was sufficient to explain the colonization outcomes observed experimentally. Here, we model *Vibrio* gut colonization in zebrafish larva by coupling stochastic colonization dynamics with bacterial population growth in the larval guts (*B_F_*) and surrounding environment (*B_W_*). Bacteria grow logistically in the host gut, experience decay in the water in the absence of hosts and are exchanged between gut and environment through colonization and expulsion. The larvae become colonized (*F_C_*) probabilistically depending on the environmental bacterial density via a Hill function, with new colonization events being determined from a binomial process. If a colonization event occurs, a number of bacteria (ξ) move from the water environment into the gut. Fish colonized with bacteria will expel a fraction of their gut bacteria back into the environment at rate σ. Most parameters were derived from experimental measurements, while a few—such as those governing colonization probability—could not be measured directly and were estimated by comparing simulations to experimental outcomes across parameter ranges (S2 Fig). A complete description of the model and parameter estimation is provided in the Methods and SI Appendix.

**Figure 3.**
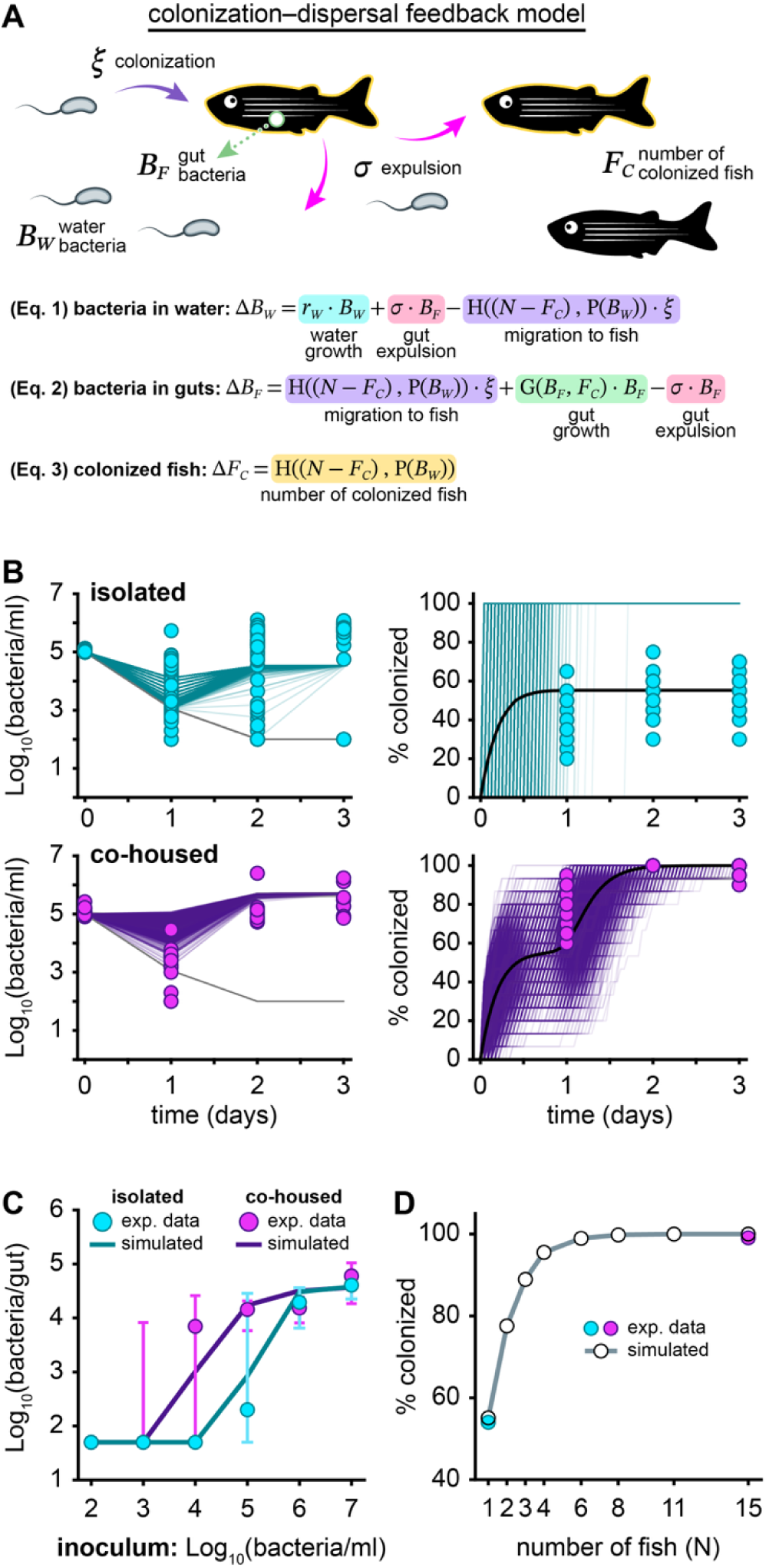
A quantitative colonization–dispersal feedback model describes rapid mass host colonization. (A) Schematic of the colonization–dispersal feedback model depicting key parameters defined in Table 1 and the equations constructed to express model dynamics. Bacterial abundance in the water (left) and the fraction of colonized animals (right) for isolated (top) and co-housed (bottom) animals over a 3 day period. Cyan and purple lines indicate model simulations in which 100% of the host population became colonized, grey lines indicate populations that did not reach complete colonization. Each line represents one of 10000 simulations (darker lines indicate overlapping lines). The average fraction of colonized individuals across simulations is indicated in black. Experimental data (derived from Fig 2B and C) are shown as cyan or purple circles for comparison. Data for the colonized population frequency was randomly subsetted to generate more points for comparison. The fraction of simulations achieving complete colonization was 54.93% and 99.92% for isolated and co-housed animals, respectively, aligning with experimental data shown in Fig 1D. (C) Gut abundances across inoculation doses for isolated and co-housed conditions. Lines show the median across 10000 model simulations after 24h. Symbols and bars indicate experimental medians and interquartile ranges (derived from Fig 1A and B). (D) Average frequency of colonized individuals after 72h as population size (*N*) increases. Each point represents the average of 10000 simulations. Model predictions align with experimental values (cyan and purple circles): for *N* = 1, the model yields 54.94% and the data yields 54% colonization; for *N*=15 the model yields 99.99% and the data yields 99% colonization frequencies. Communities of ≥4 animals approach near-universal colonization.

**Table 1.** Definitions of model parameters, variables, and functions.

| Variable or Parameter | Definition | Units |
| --- | --- | --- |
| $B_W(0)$ | Initial condition of bacteria concentration in water | bacteria/mL |
| $B_F(0)$ | Initial condition of bacteria concentration in fish guts | bacteria/mL |
| $F_C(0)$ | Initial number of fish colonized with bacteria | - |
| $N$ | Total number of fish | - |
| $r_W$ | Growth rate of bacteria in water | 1/h |
| $r_F$ | Growth rate of bacteria in fish guts | 1/h |
| $K$ | Carrying capacity of bacteria in fish guts | bacteria/mL |
| $\sigma$ | Rate of expulsion | 1/h |
| $\xi$ | Number of cells colonizing gut from water | bacteria/mL |
| $A$ | Probability function half-maximal constant | - |
| $B$ | Probability function Hill coefficient | - |
| $P(B_W)$ | Probability of colonization event occurring given density of water bacteria | 1/h |
| $H((N-F_C), P(B_W))$ | Number of uncolonized fish to become colonized | - |
| $G(B_F, F_C)$ | Growth rate of bacteria in fish guts | - |

Simulations of this model reproduced the major trends observed experimentally. In both isolated and co-housed conditions, the model captured the characteristic dip and recovery of bacterial abundance in the water and overall host colonization frequencies (Fig 3B). Colonization events occurred primarily within the first 48h, consistent with experimental observations (Fig 1D). The model also recapitulated differences in gut bacterial abundance between isolated and co-housed animals at intermediate inoculum densities and predicted colonization frequencies that closely matched experimental data (approximately 55% for isolated animals and nearly 100% for co-housed animals) (Fig 3C,D; Fig 1A,B). One limitation is that the model does not capture the high variability in *Vibrio* decay rates in the water of uncolonized fish, since we used a single-valued parameter for simplicity. Despite this, the model successfully reproduces all other major trends, indicating that it integrates sufficient mechanisms to explain observed colonization dynamics.

After validating that our model captures the essential mechanisms of colonization, we next used it to explore how connectivity through a shared water environment influences colonization outcomes as host population size increases. Simulations revealed that adding more hosts rapidly increases colonization frequency, with only four animals being sufficient to achieve an average colonization rate exceeding 94% (Fig 3D). This effect arises because each colonized host replenishes water bacterial abundance, creating a positive feedback loop that accelerates colonization of remaining hosts. In contrast, isolated animals lack this connectivity and remain dependent on the initial inoculum and *Vibrio* decay rate, making colonization less likely. These findings demonstrate that environmental connectivity among hosts is a major factor shaping colonization dynamics, transforming early probabilistic colonization events into near-complete host population colonization.

We next probed host factors that could potentially modulate the strength of the colonization–dispersal feedback loop. Simulations revealed that for isolated animals, colonization frequency depends primarily on water bacterial abundance and decay rate, with little sensitivity to host traits such as carrying capacity or expulsion rate (Fig 4A, left). In contrast, co-housed animals are strongly influenced by these parameters because they control the transfer of bacteria from the gut back into the water (Fig 4A, right). When expulsion rates are moderate, bacteria accumulate in the gut and are periodically released into the environment, sustaining water bacterial abundance and promoting colonization of other hosts. However, when expulsion is too high, bacteria are removed from the gut faster than they can grow, preventing gut populations from reaching carrying capacity and limiting environmental replenishment. Conversely, higher carrying capacities allow gut populations to reach larger sizes, increasing the amount of bacteria that are expelled and accelerating colonization of remaining hosts. Simulations illustrating the impact of all pairwise model parameters on colonization are shown in S2B Fig for both isolated and co-housed conditions. Together, these simulations reveal how host traits amplify the feedback loop and govern colonization outcomes among connected hosts.

**Figure 4.**
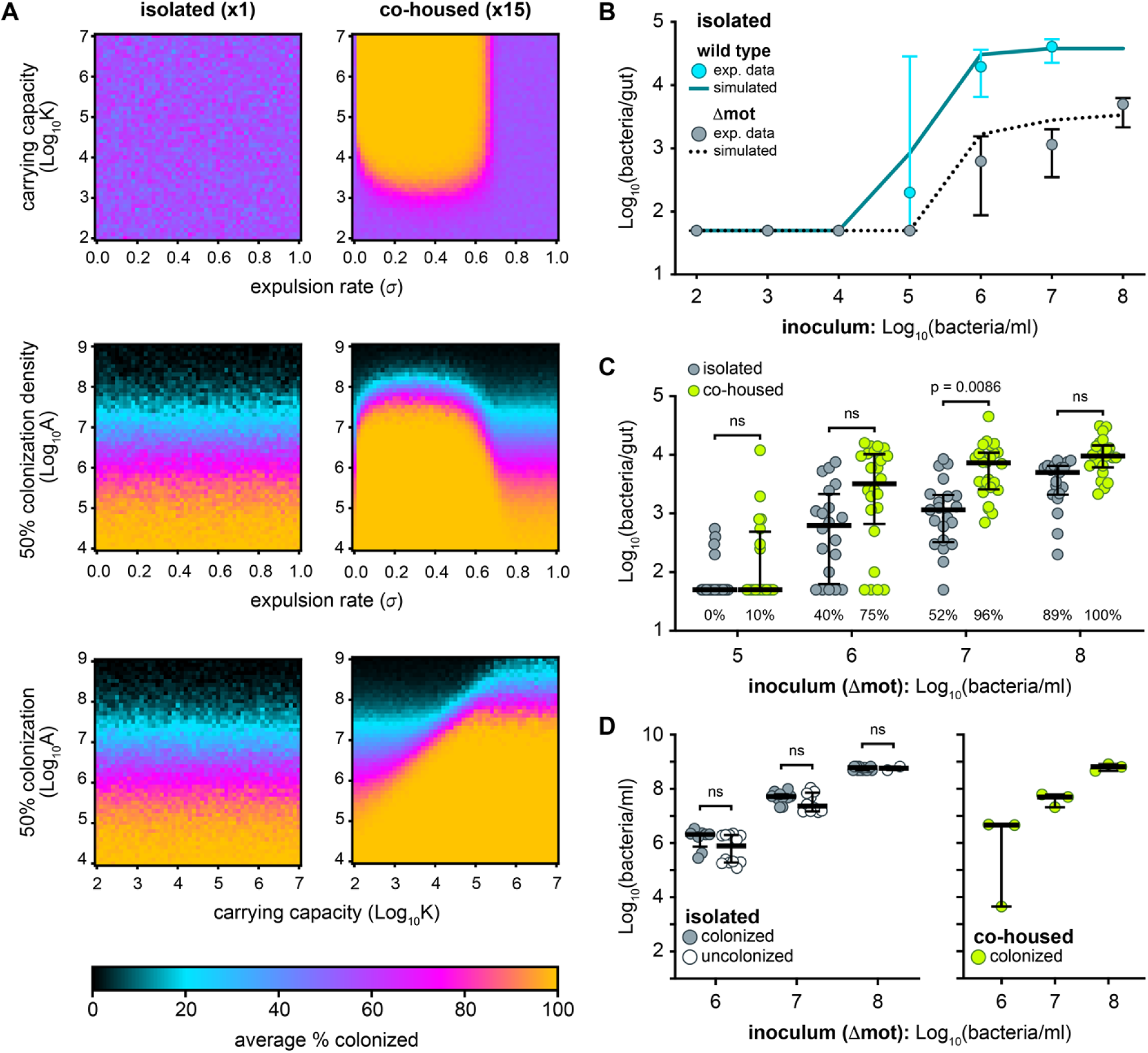
Dispersal interacts with host and bacterial traits to shape colonization thresholds. (A) Effects of carrying capacity (*K*), expulsion rate (*σ*), and the environmental density conferring 50% colonization odds (*A*) under isolated (left) and co-housed (right) conditions. Each parameter span includes 51 evenly spaced values; 100 simulations were averaged per pair. In isolated animals, colonization depends primarily on *A*; in co-housed animals, higher *K* and low-to-moderate *σ* further increase colonization. (B) Gut colonization of isolated animals inoculated with a dose range of wild-type *Vibrio* or Δmot. Lines depict simulated data; symbols and error bars indicate medians and interquartile ranges of experimental data (wild-type data derived from Fig 1A). Sample sizes for Δmot (left to right): 26, 13, 28, 27, 20, 21, 19. Statistical significance was assessed by Mann–Whitney tests. (C) Δmot gut abundances at 1 dpi in isolated and co-housed animals across an inoculum range. Sample sizes (left to right): 27, 20, 20, 24, 21, 24, 19, 22. Percentage of animals colonized (≥10^3^ bacteria/gut) is shown below each scatter plot. (D) Water abundances for the Δmot mutant in isolated (left) and co-housed (right) animals at 1 dpi. Statistical significance assessed by Mann–Whitney tests.

Finally, we asked whether the model could predict outcomes for *Vibrio* strains with altered dispersal traits. For this, we examined a swimming motility-deficient *Vibrio* mutant (Δmot), which exhibits a lower gut carrying capacity and higher expulsion rates [21]. Simulations predicted that a Δmot mutant would have a lower frequency of colonization (Fig 4B). To model the Δmot mutant, parameters for expulsion and the colonization probability function were explored to find values that aligned with the Δmot mutant colonization frequency data at an inoculum of 10^5^ bacteria/mL (S2C Fig). Further simulations using those parameters shows a shift in the colonization frequency curve toward higher inoculum densities compared to wild-type, which was confirmed by experimental data (Fig 4B,C). In isolated animals, the CD50 for the Δmot mutant is 10–100-fold higher than wild-type, while in co-housed animals, it could be partially rescued (Fig 4C). Notably, for isolated animals, water abundances of the Δmot mutant were less predictive of colonization success; even high water densities did not consistently result in gut establishment, unlike wild-type *Vibrio* where water levels strongly correlated with colonization (Fig 4D). These findings highlight that both dose and bacterial swimming motility interact to shape *Vibrio*’s colonization probability.

### Interhost transmission buffers gut bacteria against antibiotic-induced collapse

Our experiments and quantitative modeling show that mass colonization of co-housed hosts begins as a probabilistic process but becomes stabilized through a colonization–dispersal feedback loop. In this loop, colonized hosts act as bacterial reservoirs that replenish environmental abundance, enabling secondary colonization events. We reasoned that similar dynamics could govern resilience during disturbance, where recolonization from environmental or interhost sources is essential for stability [42,43]. In other words, the persistence of gut communities may depend on the same feedback mechanisms that drive initial colonization. Therefore, because zebrafish continuously shed bacteria into the water, we hypothesized that ongoing interhost transmission could buffer gut communities against antibiotic-induced collapse.

We first determined whether interhost transmission persists after initial colonization. To test this, we co-housed zebrafish harboring established gut populations of *Vibrio* strains marked with different fluorescent tags. By 24h, most animals contained small but detectable levels of transmitted bacteria, indicating continuous low-level transmission among hosts that could enable recolonization (Fig 5A).

**Figure 5:**
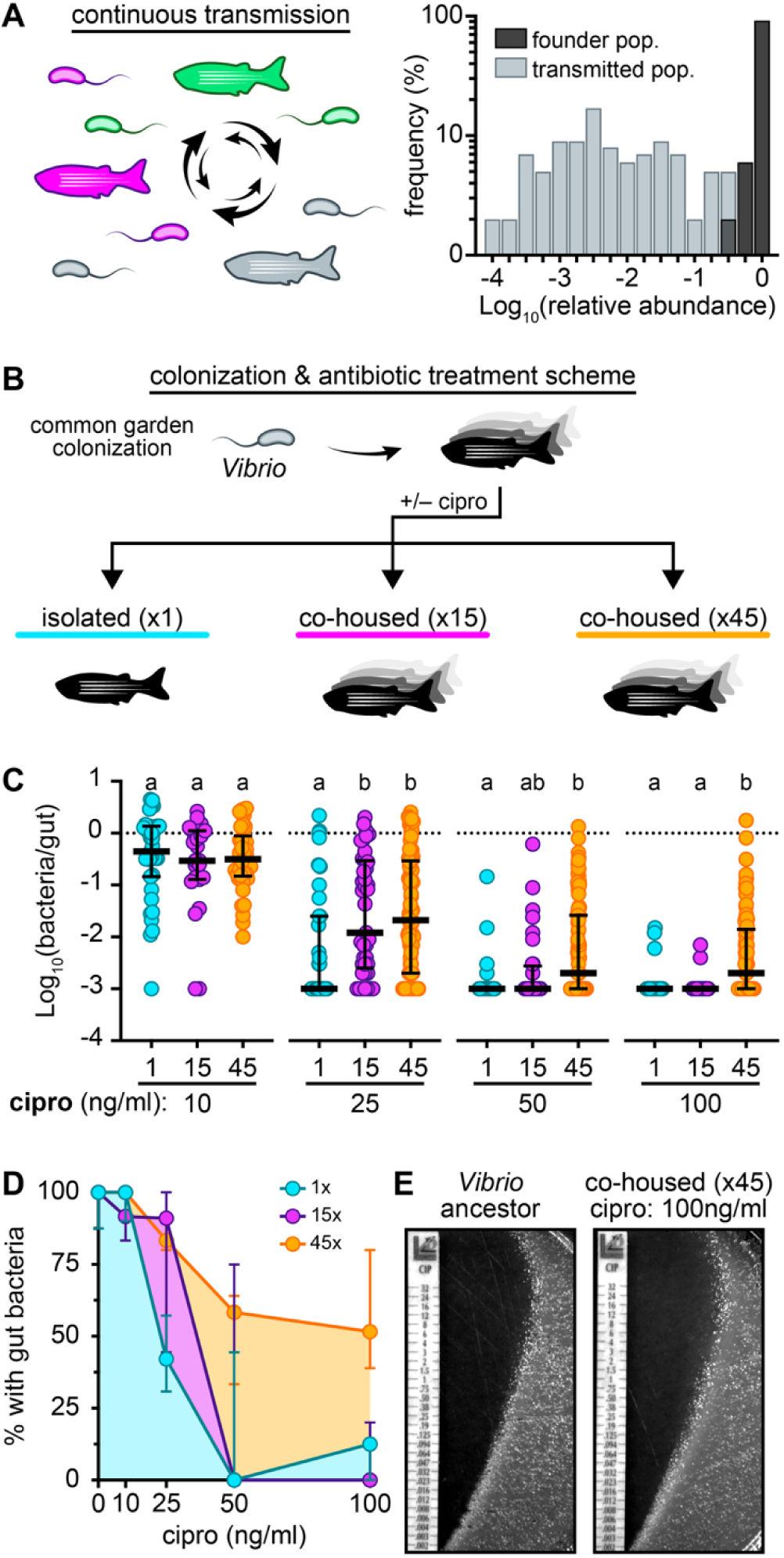
Interhost transmission buffers gut bacteria against antibiotic-induced collapse. (A) Left: Schematic of the experiment used to detect continuous interhost transmission by co-housing animals carrying differentially tagged *Vibrio* gut populations. Right: Distribution of combined relative frequencies for founder and transmitted gut populations. Sample size = 50 animals. (B) Colonization and antibiotic-treatment scheme for testing the influence of dispersal regimes on gut bacterial stability. (C) *Vibrio* gut abundances following ciprofloxacin treatment. Abundances are normalized to median abundances of untreated but similarly housed control animals (dotted lines). Letters denote statistical groupings within each antibiotic concentration as determined by Kruskal–Wallis and Dunn’s multiple comparisons tests (10 ng/mL: *p* = 0.586; 25 ng/mL: *p* = 0.0014; 50 ng/mL: *p* = 0.0019; 100 ng/mL: *p* < 0.0001). Sample sizes = 22–97 per condition. Bars indicate medians and interquartile ranges. (D) Percentage of fish with detectable gut bacteria following ciprofloxacin treatment. Lines correspond to group sizes (1, 15, 45); area under the curve is shaded for each line to visualize differences. Symbols and bars denote means and standard deviations. (E) Representative ciprofloxacin sensitivity assay comparing the ancestral *Vibrio* strain with a *Vibrio* isolate recovered from co-housed animals (group size = 45) treated with 100 ng/mL ciprofloxacin. The minimal inhibitory concentration (MIC) for all isolates, including the ancestor, was estimated to be 0.002 µg/mL.

We next tested whether co-housing enhances *Vibrio* resilience during antibiotic stress. Zebrafish were colonized for 24h under a common garden regime to establish high gut abundances, then transferred to sterile media and housed either in isolation or in groups of 15 or 45 (Fig 5B). Animals were then exposed for 24h to a concentration range of ciprofloxacin (cipro), a bactericidal fluoroquinolone antibiotic that inhibits DNA gyrase. Cipro was selected because our previous work showed that even sublethal doses of this antibiotic induce physiological alterations in *Vibrio* populations that render them susceptible to intestinal expulsion [41]. Based on our colonization–dispersal model, we therefore anticipated that perturbing gut communities with ciprofloxacin would produce outcomes highly sensitive to microbiome connectivity. Low-dose ciprofloxacin (10 ng/mL) had minimal and equivalent effects across all conditions (Fig 5C). At higher concentrations (25–100 ng/mL), *Vibrio* abundances declined sharply in all groups, but isolated hosts experienced the most severe collapses—up to 1,000-fold reductions that pushed gut populations near extinction—whereas co-housed hosts retained higher abundances and a greater fraction of colonized individuals (Fig 5C,D). For instance, at 100 ng/mL, 3% of isolated hosts retained *Vibrio* compared to 56% of animals in groups of 45 (Fig 5D). Notably, we did not detect cipro-resistant *Vibrio* isolates in co-housed conditions, ruling out resistance as the sole basis for increased stability (Fig 5E). In addition, water abundances at the onset of antibiotic exposure were equivalent across all group sizes and treatment, indicating that subsequent differences in gut population stability are not attributable to baseline variation in environmental bacterial load (S3 Fig). These results demonstrate that co-housing and interhost transmission strongly stabilize gut bacterial populations during antibiotic stress, and that this buffering effect scales, to some degree, with group size.

### Dispersal overrides antibiotics to stabilize complex gut microbiomes

Our experiments so far have focused on a single model strain of *Vibrio* in mono-association, which enabled us to dissect colonization dynamics and stability in a controlled setting. We therefore asked whether the patterns revealed by this minimal system also emerge in complex, naturally assembled consortia. To address this question, four-day-old conventionally raised zebrafish were divided into isolated or co-housed groups for 24h to allow microbiome colonization and assembly. Animals were then either treated with cipro (10 ng/mL) or left untreated for an additional 24h (Fig 6A). After treatment, gut and water samples were collected to profile bacterial communities under each housing and treatment condition (S4 Fig).

**Figure 6.**
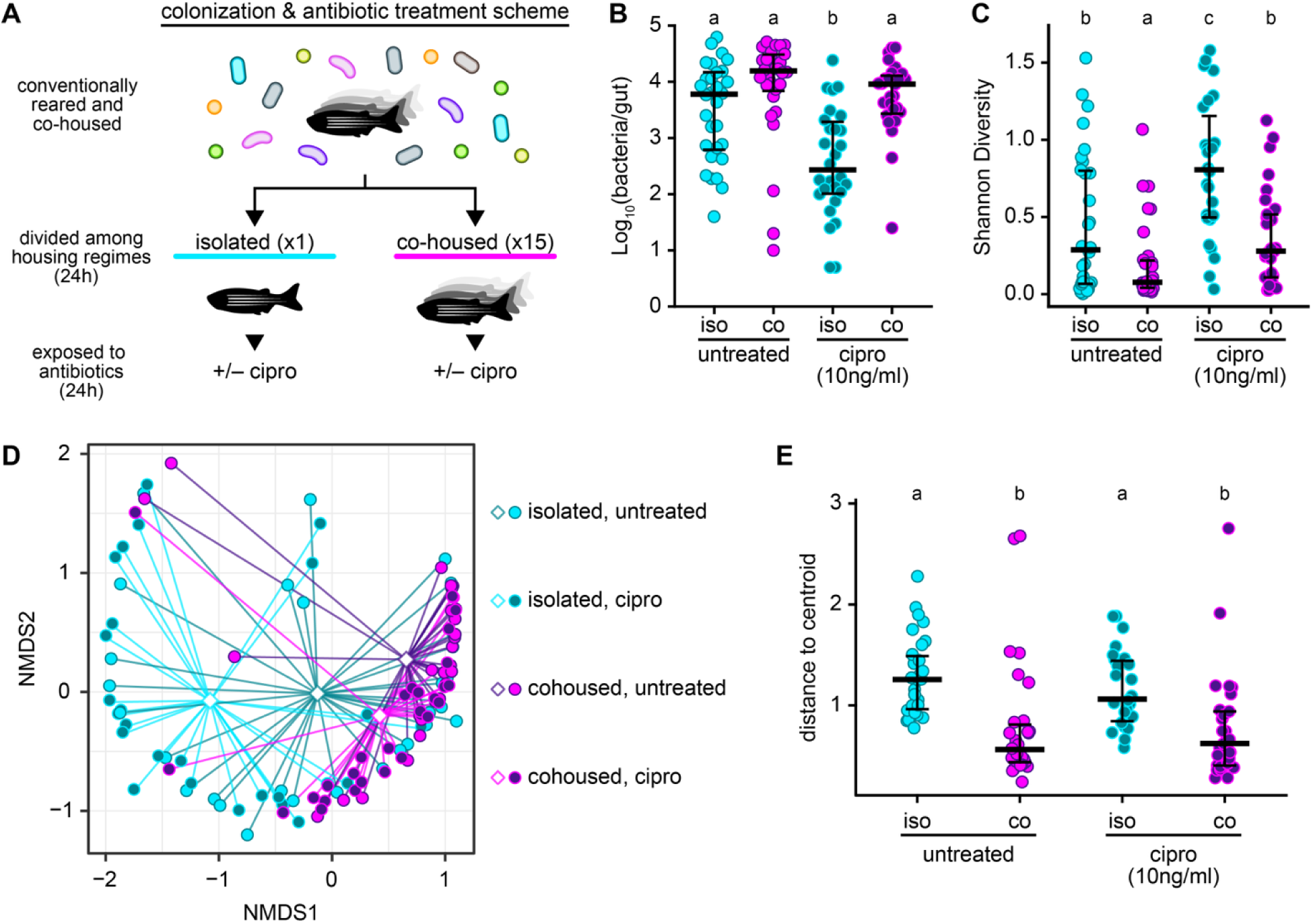
Dispersal overrides antibiotics to stabilize complex gut microbiomes. (A) Experimental scheme for colonization and antibiotic treatment. (B) Absolute gut bacterial abundances for all housing and treatment conditions (iso = isolated; co = co-housed). Bars indicate medians and interquartile ranges. Letters denote statistical groupings determined by Kruskal–Wallis and Dunn’s multiple comparisons test (*p* <0.0001). Sample sizes from left to right: 30,28,30, 29. (C) Shannon diversity index across housing and treatment groups. Bars indicate medians and interquartile ranges. Letters denote statistical groupings determined by Kruskal–Wallis test (*p* <0.0001) followed by a post-hoc pairwise Wilcoxon rank-sum test with FDR correction. (D) NMDS ordination of Bray–Curtis dissimilarities showing differences in microbial community structure across conditions. Circles denote individual samples, while diamonds indicate the centroid of each group. Connecting lines show the distance of each sample from its corresponding centroid. The NMDS stress value (stress = 0.06). (E) Distance to centroid for each condition reflects within-group community variability. Bars indicate median and interquartile ranges. Letters denote statistical groupings as determined by Kruskal–Wallis (*p* < 0.0001), with pairwise Wilcoxon post hoc comparisons (FDR-corrected).

Because larval zebrafish microbiomes are dominated by Pseudomonadota (Proteobacteria), members can be readily cultivated and visually distinguished on chromogenic media (which stains colonies based on their metabolic activity)[38,44]. This strategy provides a straightforward, scalable method for tracking community composition. Nine common colony morphologies could be classified based on color pattern and size, and representative isolates were identified by 16S rRNA sequencing (S4A Fig).

We first observed that absolute gut bacterial abundances of naturally assembled microbiomes mirrored patterns observed with *Vibrio* mono-associations. Isolated, cipro-treated animals exhibited more than a ten-fold reduction in gut bacterial load compared to untreated controls (Fig 6B). In contrast, co-housed animals maintain abundances comparable to untreated groups despite antibiotic exposure.

Assessment of alpha (within-host) diversity using the Shannon index revealed that untreated isolated and co-housed animals exhibited modest yet significantly divergent diversity levels, with co-housed animals showing lower diversity (Fig 6C). Alpha diversity increased in both housing conditions following cipro treatment, but the effect was more pronounced in isolated animals, which displayed significantly higher indices for Shannon diversity compared to co-housed animals (Fig 6C). Lower diversity in co-housed animals likely reflects greater community homogenization driven by interhost transmission, promoting convergence on a shared set of dominant taxa. In contrast, isolation combined with antibiotic treatment appears to shift microbiomes into different states due to probabilistic colonization precluding the establishment and numeric dominance of a single taxon.

This gradient of stochasticity to a more deterministic state is clear by looking at differences in taxonomic composition across larvae. Beta (between-host) diversity analysis based on Bray-Curtis dissimilarities showed that co-housed animals clustered tightly regardless of antibiotic treatment, whereas isolated animals exhibited greater inter-individual variability (Fig 6D,E). Multivariate dispersion differed significantly between housing conditions (*p* = 0.001), indicating that housing strongly influences within-group variability. In contrast, antibiotic treatment alone did not significantly affect dispersion. Further analysis revealed significant heterogeneity among housing × antibiotic groups (*p* = 0.001), suggesting that housing context modulates microbiome responses to antibiotic exposure and that interhost transmission among co-housed animals, buffers against disturbance. Consistent with these patterns, compositional profiles showed that co-housed microbiomes remained dominated by a single morphology (M1), representing multiple *Vibrio* lineages, whereas isolated fish exhibited depletion of M1 and opportunistic expansion of alternative taxa represented by M2 and M5 morphologies (representing *Shewanella*, *Pseudomonas*, *Ectopseudomonas*, *Brevundimonas*, and *Chryseobacterium* species) (S4B–D Fig).

Interestingly, although the M1 morphology was the most dominant, representative isolates from this class were also the most sensitive to cipro (S1 Table). In contrast, isolates belonging to M5 and M9 morphologies displayed substantially higher levels of cipro resistance yet failed to expand in co-housed, cipro-treated conditions (S1 Table). Therefore, persistence of the M1 morphology in treated co-housed animals likely reflects rescue through dispersal rather than intrinsic or evolved resistance.

Analysis of water samples revealed that abundances of the most predominant environmental morphologies (M1–M3) did not necessarily correlate with gut compositional shifts, in contrast to *Vibrio* mono-associations where water levels reliably predict gut outcomes. This result is still consistent with interhost dispersal shaping complex gut communities but emphasizes a role for additional layers of microbial interactions beyond bacterial abundances in different compartments (S4E–G Fig). Together, these findings demonstrate that co-housing—and by extension, gut bacterial dispersal—can override antibiotic effects, while isolation amplifies probabilistic, drift-driven divergence among hosts.

## Discussion

Our study uncovers a mechanistic bridge between the inherently stochastic process of gut bacterial colonization in individual hosts and the widespread colonization that can emerge among hosts connected by dispersal. In isolated gnotobiotic larval zebrafish, gut colonization follows a classic dose-dependent, probabilistic curve mirroring that described in invertebrate models (Fig 1A). However, we find that a bacterial dose leading to colonization half of the time in isolated hosts gives rise to mass colonization across a group (Fig 1B). We capture this transition in outcomes with a quantitative model (Fig 3), revealing that it is driven by a colonization–dispersal feedback in which early colonized hosts act as reservoirs that continuously reseed the shared environment, elevating colonization probabilities for all remaining hosts. In this way, we demonstrate how interhost dispersal can overwhelm the local filtering mechanisms and bottlenecks that ordinarily constrain colonization, fundamentally shifting the rules of assembly across scales and stages of community development.

The transition from individual stochasticity to group-level predictability for gut colonization echoes recent work showing that complex microbial systems can converge on simple, reproducible patterns at coarse scales despite underlying heterogeneity [45]. This so-called “emergent simplicity” was shown to arise in synthetic communities when metabolic cross-feeding and functional redundancy stabilize interactions and channel diverse strains toward similar, higher-level assemblages. In our system, an analogous form of collective organization emerges not from metabolic exchange but from microbiome connectivity. Once colonization occurs in even a single host, interhost transmission ignites a cascade that drives colonization and assembly toward a shared, population-level outcome. Our results therefore extend the emergent simplicity concept by showing that dispersal and transmission can act to canalize colonization outcomes. Moreover, our work provides empirical evidence illustrating how dispersal does not always behave as a neutral or noisy process but that it can be a primary driver of when, where, and how gut communities assemble.

Crucially, probabilistic colonization was required for our quantitative model to accurately describe experimental results. A simple deterministic model could not recover both the frequency of colonization in isolated and co-housed fish. This highlights the importance of stochasticity in accurately describing certain ecological dynamics, as well as elucidating the biological significance of probabilistic colonization in the assembly of gut microbiomes [9,35,36]. The dynamics of colonization and community assembly in fish depended on the stochasticity of early colonization events, as well as transmission dynamics that led to near-complete colonization in large groups. This initial stochasticity could possibly amplify priority effects, potentially leading to alternative states of microbiome composition in different metacommunities. Our study ultimately establishes a system for evaluating the interplay between competition, environmental selection, and interhost dispersal during microbiome assembly and recovery from perturbation.

Establishing *Vibrio*’s CD50 was central to experimentally resolving how dispersal tips the balance from probabilistic to mass colonization. We formalize the CD50 as an analog of the well-established infectious and lethal dose thresholds (ID50/LD50), which have been used for decades to define host filtering mechanisms and pathogen traits governing infection [46–48]. In our system, the CD50 served a similar function by providing a tractable metric for identifying factors that influence gut colonization outcomes. From an ecological perspective, associating hosts and bacteria at the CD50 places the system in a probabilistic regime in which colonization outcomes become hypersensitive to even subtle changes in niche-based or neutral processes. We demonstrated this sensitivity by showing that facilitating interhost transmission via co-housing lowered *Vibrio*’s CD50 by roughly an order of magnitude relative to isolated hosts, whereas disrupting *Vibrio*’s flagellar motility—which we previously showed is critical for resisting the expulsive mechanics of the gut—raised the CD50 by nearly two orders of magnitude (Fig 1B, Fig 4C). Similar links between motility-dependent bottlenecks and interhost transmission have been recently demonstrated in infant mouse models [49]. Together, these results highlight how dispersal and microbial traits interact to determine the CD50 curve, and how these interactions could either conceal or reveal shifts in colonization thresholds that would otherwise remain obscure.

More generally, our studies using single-member and complex communities—combined with the insights that emerge from our colonization–dispersal model—illustrate how dispersal and probabilistic colonization may interact in natural microbiomes. Members of an assembling community are expected to span a spectrum of inherent CD50 relationships with their host, determined by their traits and compatibility with host filtering mechanisms. Interhost transmission and the degree of microbiome connectivity can then shift these relationships upward or downward. Experimentally, we anticipate that systematically assessing strain-specific host colonization thresholds in isolated and co-housed conditions will uncover mechanisms that were previously masked by long-standing protocols originally designed to achieve uniform colonization via high inoculating doses.

Interpreting colonization thresholds in natural contexts will require moving beyond single host–strain relationships and recognizing that many host–microbe interactions transform gut physiology in ways that can collaterally alter the CD50s of other community members. For example, several enteric pathogens, including *Vibrio* and *Salmonella* species, modulate intestinal contractions to promote their own colonization and dissemination [50,51]. The consequences of such perturbations can be readily explored using our colonization–dispersal model, which predicts that increased expulsion may sensitize certain taxa to extinction unless they are rescued by sufficient connectivity to surrounding microbiomes. Notably, the *Vibrio cholerae* strain used in our study can amplify intestinal contractions through its Type VI Secretion System, which was previously shown to also mediate competitive exclusion of other gut bacteria [39,52,53]. Together, our work—which builds upon foundational studies in invertebrate models—sets the stage for exploring how single host–microbe interactions scale with microbiome connectivity to collectively shape microbiome assembly.

The assembly and stability dynamics revealed by our study show striking yet intuitive parallels between gut bacterial colonization and long-standing concepts in metapopulation and infectious-disease ecology. Classical host–pathogen frameworks predict that “rescue effects” arise when extinction risk declines as the number of occupied host patches increases across a spatially structured population [43,54–56]. Similar rescue-type effects have been theoretically anticipated in modified SIR (Susceptible, Infective, Recovered) models where hosts are indirectly connected through shared environmental reservoirs [57,58]. Although we did not explicitly model direct host-to-host transmission, we observed an analogous rescue effect in which increasing host number—and thus effectively increasing the number of host patches—reduces bacterial extinction frequency via environmental connectivity alone. While rescue effects have been widely documented in microbial systems, they are typically attributed to processes operating within populations, such as mutation or standing genetic variation [59,60]. Empirical evidence for rescue effects mediated purely by indirect connectivity, however, remains limited. Here, we demonstrated—both empirically and theoretically—that such rescue dynamics can arise even when connectivity between hosts or patches occurs solely through a shared environment. Although resident gut microbes are typically considered distinct from pathogens, our results suggest two complementary interpretations: (i) the processes governing the transmission and persistence of gut bacteria share fundamental principles with pathogen ecology, and (ii) mechanistic insights from gut bacterial colonization dynamics can feed back to inform our understanding of pathogen spread. In this way, our framework for predicting gut colonization probabilities may ultimately provide a useful bridge between microbiome assembly and classical disease-transmission theory.

The use of gnotobiotic zebrafish provided essential experimental tractability for investigating dispersal and gut colonization, enabling direct measurement of bacterial abundances across both hosts and their surrounding environment under controlled housing configurations. Although our system is aquatic, the principles that emerged from this work likely generalize in terrestrial hosts, including humans [11,26,61,62]. The parameters that govern colonization in larval zebrafish—bacterial persistence in water, movement between environmental reservoirs, intestinal growth and expulsion, and the frequency of host exposure—have clear analogs in natural and built environments [26,61,63,64]. Human habitats across urban and rural settings contain diverse microbial landscapes in which bacteria persist on surfaces, in the air, and within shared spaces, collectively forming the transmission networks that define colonization–dispersal dynamics [63,64]. Notably, a key host trait not captured in our study, however, is sociality. We studied zebrafish larvae prior to the onset of social behaviors such as shoaling, yet in many animals social interactions serve as corridors for microbial transmission, inspiring the concept of the “social microbiome” [65–69] In humans, patterns of microbiome composition show that socially connected individuals—from cohabiting family members to residents of Honduran villages—share more gut strains than those with minimal interaction, a trend we surmise reflects a reduction in colonization thresholds for socially acquired taxa [29,70–72].

Finally, insights from our study may also have implications for microbiome therapies. The engraftment of beneficial microbes, maintenance of community stability, and recovery from perturbations such as antibiotic treatment depend not only on microbial traits but also on the dispersal regimes hosts are embedded in [73–77]. Our results illustrate how isolated hosts become prone to drift and colonization failure, whereas co-housing or shared environmental exposure can reinforce dispersal pathways and enhance resilience [74,75,78,79]. Indeed, it may be informative to re-examine many classic microbiome studies through the lens of microbiome connectivity. Together, these observations help reimagine transmission networks as powerful levers for shaping microbiomes. More broadly, our findings emphasize that harnessing dispersal—alongside niche-based processes—should be central to strategies aimed at preserving the integrity and function of the dense, complex, and interconnected microbial ecosystems that impact host health.

## Supplemental Figures

**S1 Figure.**
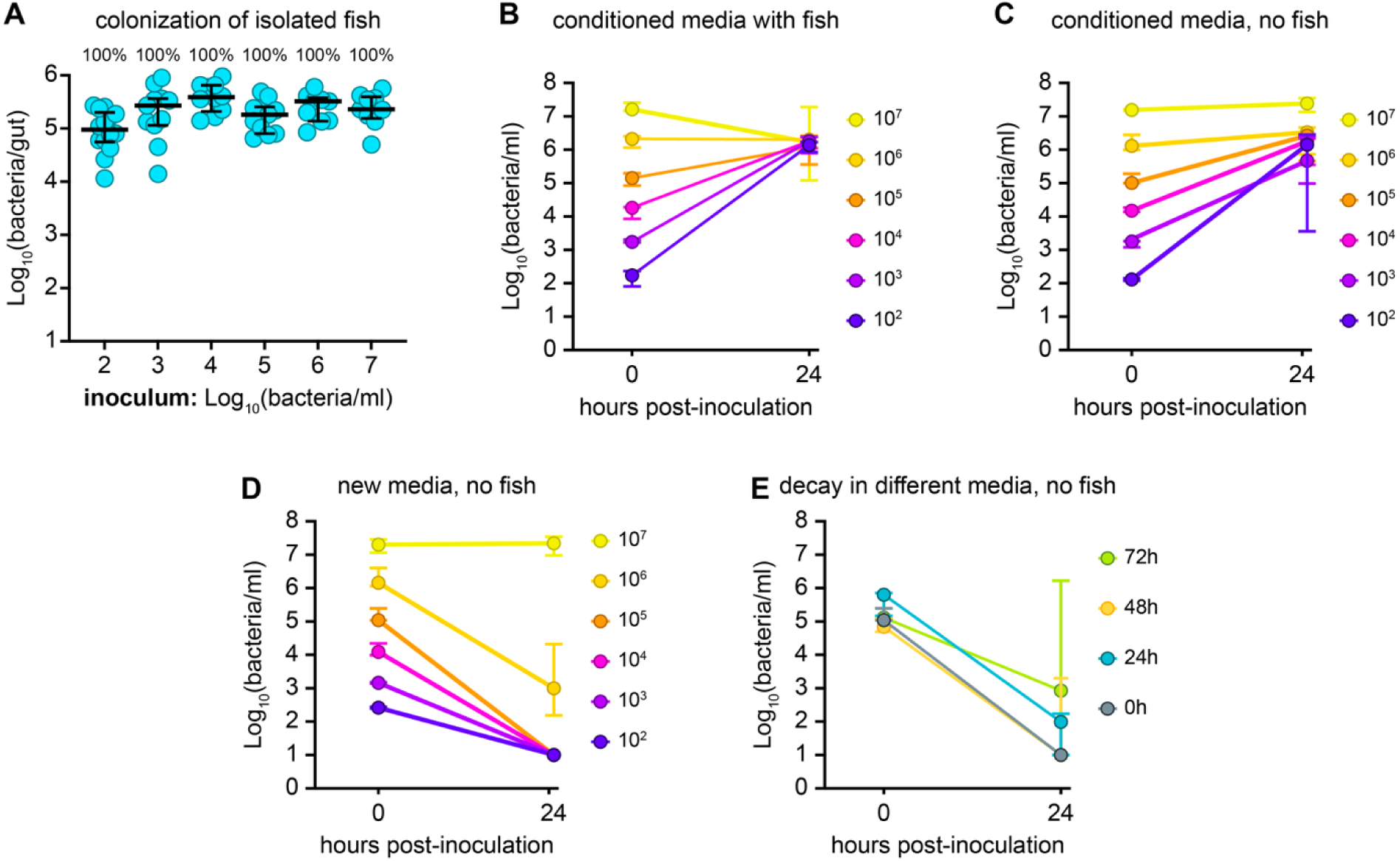
Host-conditioned water increases *Vibrio* persistence and masks dose-dependent colonization dynamics. (A) *Vibrio* gut abundances at 24h post-inoculation (hpi) for isolated fish inoculated in conditioned embryo media (EM). Bars indicate medians and interquartile ranges. Sample sizes (left to right): 14, 11, 10, 11, 9, 9. Percentages reflect the fraction of colonized animals (containing ≥ 10^3^ bacteria/gut) for each inoculation condition. (B) Initial and final *Vibrio* water abundances from the experiment shown in panel A, sampled at 0 and 24 hpi. Lines depict the trajectory of water abundances at each inoculation dose. Symbols and bars indicate medians and ranges. (C) Plot depicts initial and final *Vibrio* water abundances as in panel B, except fish were removed from EM prior to inoculation. Samples were collected at 0 and 24 hpi. Lines depict water trajectories at each dose; symbols and bars indicate median and ranges across 3 biological replicates. (D) *Vibrio* viability decay trajectories in sterile, unconditioned EM. Samples collected at 0 and 24 hpi. Lines depict water trajectories at each dose; symbols and bars indicate median and ranges across 3 biological replicates. (E) *Vibrio* viability in EM conditioned by fish for 0, 24, 48, or 72h prior to inoculation with 10^5^ bacteria/mL. Samples collected at 0 and 24 hpi. Lines depict water trajectories for each conditioning duration; symbols and bars indicate medians and ranges across 3 biological replicates.

**S2 Figure.**
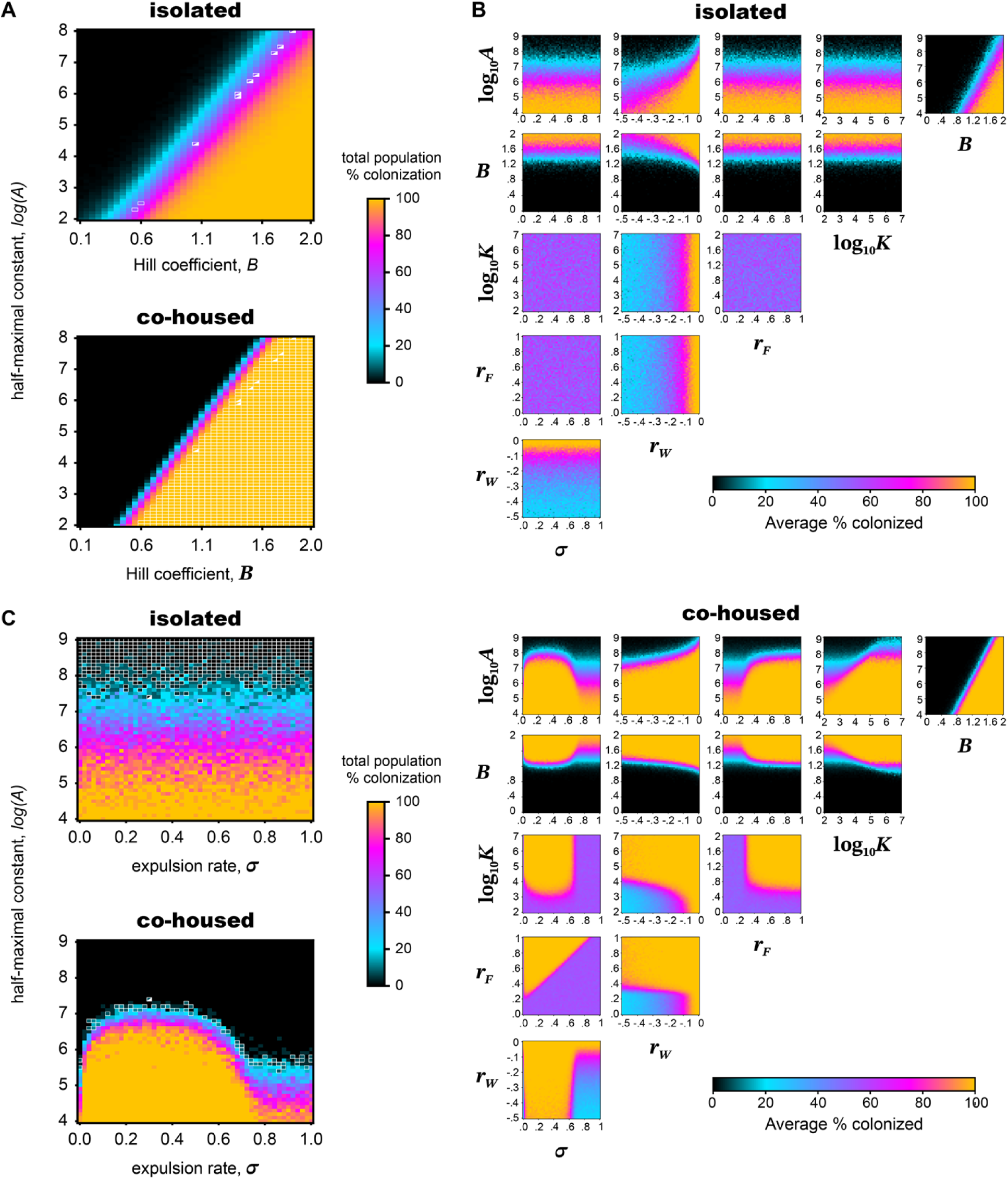
Colonization–dispersal model simulations. (A) Average frequency of simulations resulting in 100% colonization for isolated (top) and co-housed (bottom) conditions used to determine probability of colonization function parameters *A* and *B*. Simulations were ran across a wide range of values, with 2500 simulations being run for each parameter pair. 61 evenly spaced values (*x*) from 2 to 8 were used for *A* = 10^x^, and 39 evenly spaced values from 0.1 to 2.0 were used for *B*. White squares indicate values that were between 53–55% and 98–100% colonized for the isolated and co-housed conditions, respectively (roughly +/- 1% of the data). Squares half-filled diagonally indicate parameter pairs in both the isolated and co-housed conditions that aligned with the data, the median value of these parameter pairs was used for simulations. (B) Simulated average frequency of colonization across pairwise parameter variations in isolated (top) and co-housed (bottom) conditions. 100 simulations were ran for each parameter pair across a wide range of parameter values. For the isolated growth condition, only parameters altering the probability of colonization function (*A* and *B*) and the decay rate of the water bacteria (*r_W_*) affect the frequency of colonization, however, under the co-housed condition, the effect of parameters such as carrying capacity (*K*) and expulsion (*σ*) becomes evident. (C) Average frequency of simulations where 100% of the population became colonized for isolated (top) and co-housed (bottom) conditions to determine the *Vibrio* motility mutant parameter values for the expulsion rate (*σ*) and 50% probability of colonization parameter *A*. Simulations were ran across a wide range of values, with 25 simulations being run for each parameter pair. 51 evenly spaced values (*x*) from 4 to 9 were used for *A* = 10^x^, and 51 even spaced values from 0.0 to 1.0 were used for *σ*. White squares indicate values that were between 0–5% and 5–15% colonized for the isolated and co-housed conditions, respectively (roughly +/- 5% of the data). Squares half-filled diagonally indicate the parameter pair in both the isolated and co-housed conditions that aligned with the data and was used for simulations.

**S3 Figure.**
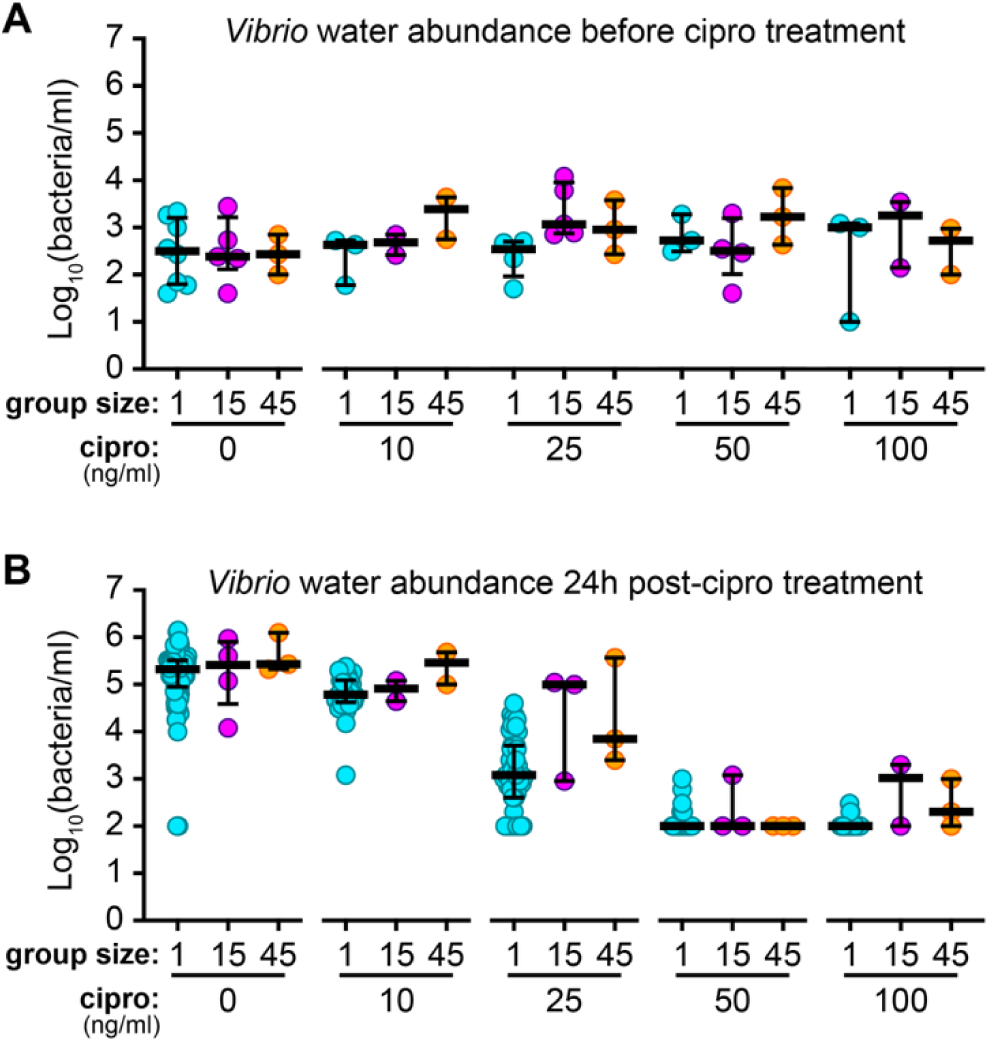
*Vibrio* Water abundances before and after ciprofloxacin treatment. (A) Water abundances in isolated and co-housed groups prior to ciprofloxacin treatment. Bars indicate medians and interquartile ranges. Sample sizes = 2–8 per condition. Kruskal–Wallis and Dunn’s multiple comparisons tests did not detect any significant differences across all water samples (*p* > 0.05). (B) Water abundances 24h after ciprofloxacin exposure. Bars indicate medians and interquartile ranges. Sample sizes = 2-84 per condition.

**S4 Figure.**
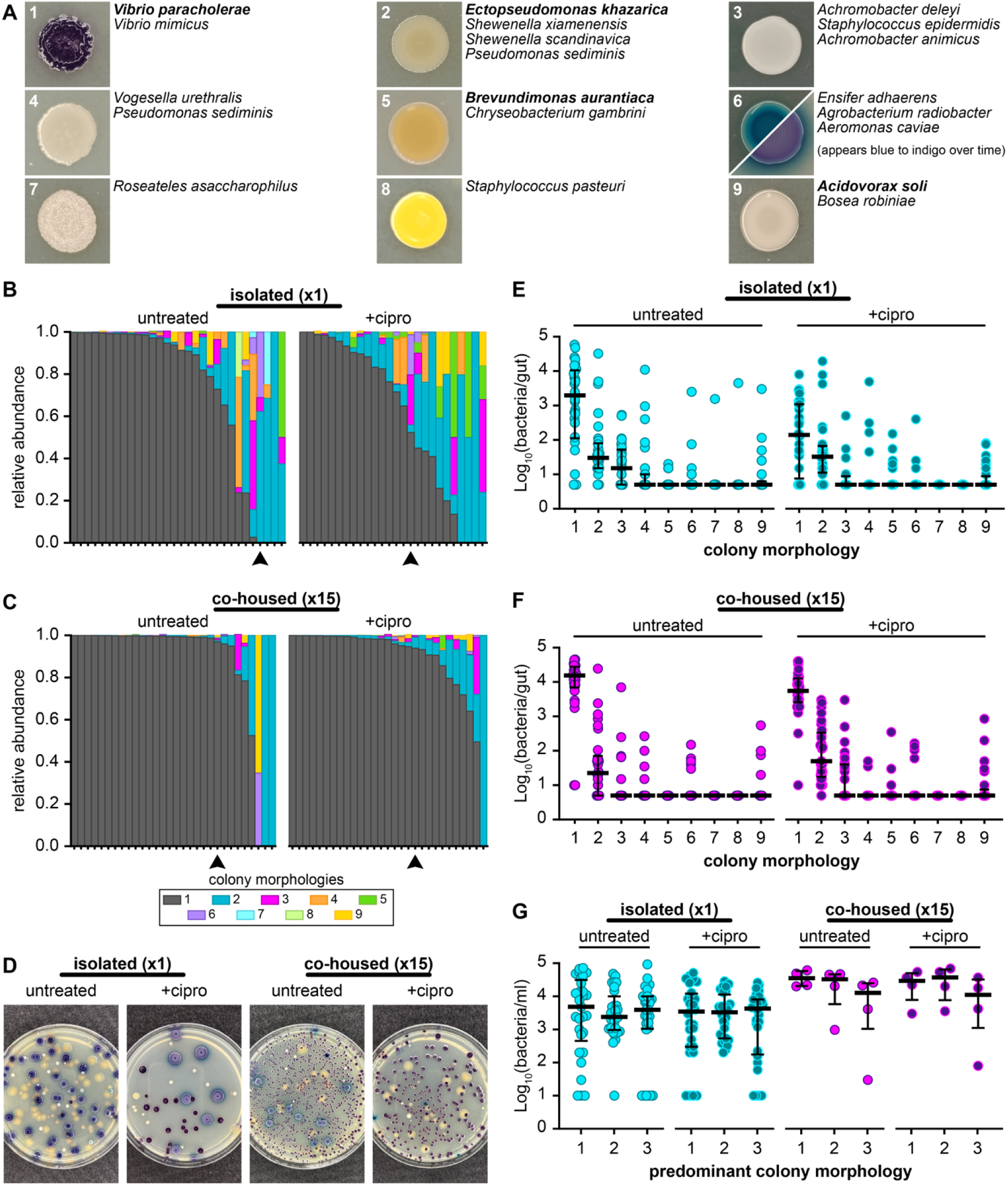
Gut microbiome profiling of conventionally reared larval zebrafish. (A) Representative images of the nine most common colony morphologies (M1–M9) isolated from conventional larval zebrafish on chromogenic media. Species associated with each morphology were identified by 16S rRNA gene sequencing; predominant lineages are shown in bold. (B) Relative abundances of M1–M9 in isolated animals. Arrowheads indicate individual fish for which corresponding plate images are shown in panel D. (C) Relative abundances of M1–M9 in co-housed animals. Arrowheads indicate individual fish for which corresponding plate images are shown in panel D. (D) Representative chromogenic agar plates for isolated and co-housed fish under untreated and ciprofloxacin-treated conditions. (E) Gut bacterial abundances of M1–M9 in isolated animals. Bars indicate medians and interquartile ranges. Sample sizes (untreated, treated): 30, 28. (F) Gut bacterial abundances of M1–M9 in co-housed animals. Bars indicate medians and interquartile ranges. Sample sizes (untreated, treated): 30, 29. (G) Bacterial water abundances of the three predominant water morphologies (M1–M3) under isolated and co-housed conditions. Bars indicate medians and interquartile ranges. Sample sizes (left to right): 29, 28, 4, 4.

**S1 Table. Bacterial strains used in this study.**

(PDF)

**S1 Data. File (.xlsx) containing all plotted numerical data.**

(XLSX)

**S2 Data. File (.txt) containing 16S rRNA sequences for bacteria isolated from conventional zebrafish.**

(TXT)

**S1 Appendix. Description of parameter estimation for the quantitative colonization–dispersal model.**

(PDF)

## Methods

### Bacterial strains and culture

*Vibrio cholerae* ZWU0020, originally isolated from the zebrafish intestine [38], was used along with fluorescently tagged derivatives. The dTomato-expressing ZWU0020 strain was generated previously by Tn7-mediated chromosomal insertion of a constitutively expressed dTomato cassette [80]. For this study, we constructed a matched mNeonGreen-expressing ZWU0020 strain using the same Tn*7* transposition workflow. Briefly, the mNeonGreen expression cassette [81] was delivered by triparental conjugation using an *E. coli* SM10 donor strain carrying the transposase-containing pTNS2 helper plasmid, and an *E. coli* SM10 donor strain carrying a pTn7xTS mNeonGreen tagging vector. Gentamicin-resistant recombinants were colony-purified, screened by fluorescence microscopy, and verified for correct insertion at the chromosomal attTn7 site by PCR using primers WP11 (5′-CACGCCCCTCTTTAATACGA-3′) and WP12 (5′-AGGGTACCGATGTTGACCAG-3′) [80]. A motility-deficient mutant (*Vibrio* Δmot), carrying a markerless, in-frame deletion of the *pomAB* locus, was previously constructed as described [21]. Archived bacterial stocks were maintained in 25% glycerol at −80 °C and inoculated directly into 5 mL tryptic soy broth (TSB) for overnight growth (∼16 h, 30 °C, shaking) prior to experiments. List of bacterial strains used in this study can be found in S1 Table.

### Ethics statement

All experiments with zebrafish were done in accordance with protocols approved by the University of California Irvine Institutional Animal Care and Use Committee (protocol AUP-23-126) and followed standard protocols [82]. Specific handling and housing of animals during experiments are described in detail below. All zebrafish used in this study were larvae, between the ages of 4- and 7-days post fertilization (dpf).

### Germ-free derivation and bacterial association

Wild-type zebrafish (AB) or zebrafish carrying the Tg(tnfα:GFP) transgene were derived germ-free following previously published protocols [83,84]. In brief, fertilized eggs from adult zebrafish mating pairs were collected and incubated in sterile embryo media (EM) containing ampicillin (100 μg/mL), gentamicin (10 μg/mL), amphotericin B (250ng/mL), tetracycline (1 μg/mL), and chloramphenicol (1 μg/mL) for approximately 6 hours. Embryos were then washed with EM containing 0.1% PVP iodine, followed by a wash with 0.003% sodium hypochlorite, and finally a wash with sterile EM. Lastly, embryos were dispensed into T75 tissue culture flasks (Genesee Scientific) containing 50mL of sterile EM at a density of 1 embryo/mL of EM and kept at 28.5 °C. For bacterial associations, overnight cultures were pelleted (1mL, 2 min, 7,000 × g) and washed twice in sterile EM before inoculation into the water column.

### Transfer of fish into non-conditioned media

At 4 dpf germ-free larvae were collected into a sterile 40µm mesh strainer and gently rinsed with sterile EM. Rinsed fish were then transferred into a petri dish containing fresh sterile EM. Visible chorion remnants were removed and unhealthy or underdeveloped animals were excluded. Media in the dish was then replaced with fresh sterile EM prior to experimental setup.

### Cultivation based measurements of abundances

Fish intestines were harvested and placed into Eppendorf tubes containing 500 µl of sterile saline (0.7% NaCl) and 100 µl of 0.55 mm zirconium oxide beads. Samples were homogenized using a NextAdvance Bullet Blender at power setting 4 for 60s. Water samples and homogenized gut samples were serially diluted in saline, plated on TSA, and incubated at 30 °C. Colony forming units (CFUs) were counted the following day. All reported abundances (bacteria/mL or bacteria/gut) represent CFUs measured on tryptic soy agar (TSA) plates. Abundance measurements reported throughout the text were pooled from a minimum of three independent experiments. Samples with zero countable colonies were assigned the limit of detection. For gut samples, the limit of detection was 50 CFU/gut for figures 1–5 and S1 Fig, and 5 CFU/gut for figure 6 and S4 Fig For water samples, the limit of detection was 100 CFU/mL for figures 1–5 and S1 Fig, and 10 CFU/mL for S4 Fig Data were plotted and analyzed using GraphPad Prism (Version 10.6.1). Statistical tests are noted in the figure legends; briefly, differences between two groups were assessed using Mann–Whitney tests, and comparisons among three or more groups were performed using Kruskal–Wallis tests with Dunn’s multiple comparisons. Statistical significance was defined as *p* < 0.05. All data values plotted in the figures can be found in the supplemental spreadsheet S1 Data.

### Dose-response colonization assays

#### Isolated fish in conditioned embryo media

Germ-free zebrafish were maintained from 0–4 days in sterile EM at a density of 1 fish/mL. At 4 dpf fish were inoculated directly in their conditioned (lived-in) media with *Vibrio* at concentrations ranging from 10^2^-10^7^ bacteria/mL. Following inoculation, individual fish along with 1 mL of inoculated conditioned media were transferred into separate wells of a sterile 48-well plate (1 fish/ml in each well). Plates were then sealed with a gas-permeable polyurethane membrane (Breath-Easy) and incubated at 28.5 °C. Water samples were collected at 0 and 24 hours post inoculation (hpi) and gut samples were collected at 24 hpi. Both gut and water samples were plated on TSA for bacterial enumeration. Colonization was defined as a gut abundance ≥10^3^ bacteria/gut.

#### Isolated vs co-housed fish in non-conditioned embryo media

Washed fish were transferred either into flasks containing 15 mL sterile EM or into petri dishes containing sterile EM. Fish in both flasks and petri dishes were maintained at the same density of 1 fish/mL of EM. Flasks and petri dishes were then inoculated with *Vibrio* across a concentration range of 10^2^-10^7^ bacteria/mL. Fish initially housed in petri dishes along with their inoculated media were then transferred into individual wells of a sterile 48-well plate (1 fish/mL in each well). Plates were then sealed with gas-permeable membranes (Breath-Easy). Both plates and flasks were incubated at 28.5 °C. Water samples were collected at 0, 1, 2, and 3 days post-inoculation (dpi), and gut samples were collected at 1, 2, and 3 dpi.

Gut and water samples were plated on TSA for bacterial enumeration. Colonization was defined as a gut abundance ≥10^3^ bacteria/gut.

### Transmission assays

#### Host bacterial shedding measurements

Flasks containing 4-day-old germ-free fish were inoculated with *Vibrio* at 10^7^ bacteria/mL and incubated at 28.5 °C for 24h to ensure colonization. At 5 dpf, fish were washed by serially transferring them through multiple wells of sterile EM to remove external bacteria, leaving only gut-associated microbes. Individual fish were then transferred into wells containing 1 mL of sterile EM. Water from each well was sampled immediately (0h) and again after 8h and 24h of incubation at 28.5 °C. Gut samples were collected at 24h. Gut and water samples were plated on TSA for bacterial enumeration. Shedding rates (Fig 2D) were determined using 8h timepoint data and were calculated as (CFU_24h_ - CFU_0h_) / 8h). For the quantitative model, expulsion rates were estimated using the 24h dataset which provided the required gut abundance measurements. Expulsion rate was defined as the fraction of gut population shed into the water every hour (shedding rate / gut abundance). Details of the expulsion rate calculation are provided in sheet ‘Figure 2D’ of the S1 Data file.

#### Transmission from colonized to germ-free zebrafish

Flasks containing 4-day-old zebrafish carrying the *Tg(tnfa:GFP)* transgene were inoculated with *Vibrio* at 10^7^ bacteria/mL and incubated at 28.5 °C for 24h to ensure colonization. At 5 dpf, colonized source fish were washed by serially transferring them through multiple wells of sterile EM to remove external bacteria, leaving only gut-associated microbes. In parallel, groups of germ-free age matched AB zebrafish were transferred into flasks of sterile, non-conditioned EM. A single washed source fish was then introduced into each flask of 14 recipient fish to maintain animal density at 1 fish/mL of media. Water samples were collected 0h and at 24h after introducing the source fish. Gut samples were collected at 24h after introduction of the source fish (6dpf). Both gut and water samples were plated on TSA for bacterial enumeration. Source fish were identified by fluorescence microscopy (Leica MZ10F stereomicroscope) based on GFP expression from the *Tg(tnfa:GFP)* transgene.

#### Transmission between colonized zebrafish

Flasks containing 4-day-old germ-free fish were inoculated with *Vibrio ZWU0020*, *Vibrio ZWU0020 attTn7::mNeonGreen*, or *Vibrio ZWU0020 attTn7::dTomato* at 10^7^ bacteria/mL and incubated at 28.5 °C for 24h to ensure colonization. At 5 dpf, fish were washed by serially transferring them through multiple wells of sterile EM to remove external bacteria, leaving only gut-associated microbes. Washed fish were then co-housed in a 1:1:1 ratio (five fish per strain) in flasks containing 15 mL sterile EM to maintain density of 1 fish/mL. After 24h of co-housing, gut and water samples were collected and plated on TSA for bacterial enumeration. Fluorescent markers were used to identify strain identity within each gut. For each animal, the most abundant gut strain was designated the founder (original colonizer), while less abundant strains were classified as transmitted or acquired bacteria.

### Antibiotic perturbation assays

#### Single-strain (gnotobiotic) perturbation across group sizes

Flasks containing 4-day-old germ-free fish were inoculated with *Vibrio* at 10^7^ bacteria/mL and incubated at 28.5 °C for 24h to ensure colonization. At 5 dpf, fish were washed by serially transferring them through multiple wells of sterile EM to remove external bacteria, leaving only gut-associated microbes. Washed fish were then transferred into sterile EM under three housing conditions: isolated individuals, groups of 15, or groups of 45. All housing conditions maintained a density of 1 fish/mL of media. Each group was then treated with ciprofloxacin (0, 10, 25, 50, or 100 ng/mL) and incubated at 28.5 °C for 24h. At 6 dpf (24 h post treatment), gut and water samples were collected and plated on TSA for bacterial enumeration. Normalized gut abundances were calculated by dividing the bacterial count of each treated fish by the median bacterial count of untreated fish of the same group size. Additionally, 6 colonies from 4 untreated fish and ten colonies from 5 ciprofloxacin-treated (100ng/mL) fish were isolated and cultured for subsequent MIC determination. All isolates were acquired from animals housed in groups of 45.

#### Complex-community (conventional) perturbation and community profiling

Zebrafish embryos were collected from adult mating pairs and transferred into a T75 tissue culture flask (Genesee Scientific) with sterile EM at a density of 1 fish/mL. At 4 dpf, fish and their media were either transferred into a 48-well plate (1 fish/mL per well) or into flasks containing 15 fish in 15 mL media. At 5 dpf, fish in both housing conditions were treated with 0 or 10 ng/mL ciprofloxacin and incubated at 28.5 °C for 24h. Gut and water samples were collected at 6 dpf (24h post treatment), plated on Universal Differential ChromoSelect Medium (Sigma-Aldrich), and incubated at 30 °C overnight for bacterial enumeration. Nine distinct morphologies were identified and used as proxies for community composition.

### Isolation and 16S rRNA sequencing of gut bacterial strains

Multiple colonies representing each distinct morphology were picked from Universal Differential ChromoSelect Medium (Sigma-Aldrich) plates and streaked for purification. Single colonies from purified streaks were grown in TSB and glycerol stocks (25%) were prepared and stored at −80 °C. Genomic DNA was extracted using the Promega Wizard Genomic DNA Purification Kit according to the manufacturer’s instructions. The 16S rRNA gene was PCR amplified using universal bacterial primers 8F (AGAGTTTGATCCTGGCTCAG) and 1492R (GGTTACCTTGTTACGACTT) [85]. Amplicons were purified using the Zymo Clean and Concentrate Kit and submitted for Sanger sequencing (Genewiz). Resulting 16S sequences were aligned and analyzed in Geneious Prime (v2026.0.2), and taxonomic identity was determined using BLAST searches against the NCBI 16S rRNA (bacteria and archaea) database. Isolates sequenced were labeled ‘C1’-‘C29’; more details can be found in S1 Table and S2 Data.

### Bacterial environmental persistence

Conditioned embryo media was generated by housing germ-free larval zebrafish in sterile EM for defined intervals. For the 24 h, 48 h, and 72 h conditioning periods, 4-day-old larvae were transferred into fresh sterile EM and allowed to condition the media for 24, 48, or 72h, respectively. For the 96h conditioning period, germ-free embryos remained continuously in their original EM from 0–4 dpf without transfer. Conditioned media was collected and transferred into sterile tubes for subsequent experiments. Overnight *Vibrio* cultures were washed twice in sterile EM by centrifugation (1 mL, 2 min, 7,000 × g) and resuspended in 1 mL EM. Washed cells were inoculated into either sterile, non-conditioned EM or conditioned EM (24h, 48h, 72h, or 96h) at final concentrations ranging from 10^2^-10^7^ bacteria/mL and mixed thoroughly by vortexing. One milliliter of each inoculated medium was dispensed into wells of a 48-well microtiter plate, sealed with a gas-permeable polyurethane membrane (Breath-Easy), and incubated at 28.5 °C for 24h. Samples were collected at 0 and 24 hpi and plated on TSA for bacterial enumeration.

### Determination of ciprofloxacin minimum inhibitory concentration

Overnight bacterial cultures were washed twice in sterile EM by centrifugation (1 mL, 2 min, 7000 × g) and resuspended in 1 mL sterile saline (0.7% NaCl). Cells were then diluted 1:1000 and lawns were prepared by spreading with a sterile cotton swab onto TSA plates. Ciprofloxacin MIC test strips were placed on the inoculated agar surface, and plates were incubated at 30 °C overnight. The following day, the MIC was recorded as the lowest concentration at which bacterial growth was inhibited. Plates were imaged using the Leica MZ10F stereomicroscope.

### Community diversity analysis

All analyses were performed in RStudio (v.2025.9.2.418) with R (v6.2.6.1) using tidyverse (v2.0.0) and vegan (v2.6.4). The Shannon diversity index (alpha diversity) for each sample was calculated using the *diversity* function. Beta-diversity was assessed using the Bray-Curtis dissimilarities and visualized with non-metric multidimensional scaling (NMDS; *metaMDS*, *K*=3, *trymax* =999). The group centroids were calculated in ordination space, and within-group variability was quantified as the distance of each sample to its group centroid. For the Shannon diversity index and Distance to Centroid, normality within groups was assessed using Shapiro tests. Because assumptions of normality were not met, differences among conditions were tested using Kruskal–Wallis tests, followed by pairwise Wilcoxon tests with false discovery rate (FDR) correction. Statistical grouping was assigned based on adjusted *p*-values (α = 0.05). Homogeneity of multivariate dispersion was assessed using *betadisper*, followed by permutation tests (999 permutations). Visualizations were generated with ggplot2 (v4.0.1), showing individual samples, medians, and interquartile ranges with compact letter display indicating statistically distinct groups.

### Model description

We consider a host-gut microbe model describing the colonization and transmission of bacteria in larval zebrafish. The bacteria can be found either in the water environment (*B_W_*) or in the zebrafish larval gut (*B_F_*). *N* represents the total number of fish and *F_C_* is the number of fish colonized with the bacteria, determined by the presence of bacteria in the gut.

The probability of a fish being colonized by the bacteria is conditional upon the density of bacteria present in the water (*B_W_*), thus, we represent this with the conditional probability function P(*B_W_*) which takes the form of the Hill function

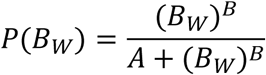

where *A* and *B* are the half-maximal constant and Hill coefficient, respectively.

To determine how many fish are colonized, we use a random binomial function to determine the number of successes (i.e. colonization events) given the number of remaining uncolonized fish (*N*-*F_I_*) and the colonization function (P(*B_W_*)). This binomial function is represented in the model as H(*N*-*F_I_,* P(*B_W_*)). To capture the dynamics of bacteria moving between the water environment and the gut, we let ξ be the number of bacteria to enter the fish gut upon colonization. Bacteria are expelled from the gut back into the water environment at rate *σ*, representing the fraction of total gut bacteria that is expelled into the water per hour.

The growth of the bacteria in the fish gut is modeled by the logistic growth function

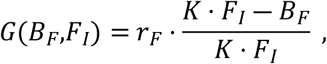

where *r*_*F*_ is the growth rate, and *K*, the carrying capacity of an individual fish, is scaled by the number of fish currently colonized. Bacteria in the water environment exhibit different growth patterns, and in fact are unable to grow in the water in the absence of fish, yielding a negative growth rate of *r*_*W*_<0. This is then captured by equations (1)-(3) in Fig 3A. A description of each parameter, variable, and function can be found in Table 1. A complete description of parameter estimation is provided in supplemental file S1 Appendix. Python version 3.13.1 was used to run all code and simulations. The code is available on Figshare (https://figshare.com/s/e553a43acd7655c53b4e).

## Acknowledgments

We are grateful to members of the Wiles lab and to Dr. Michael Parsons (University of California, Irvine) for their support in establishing and maintaining our zebrafish colony. We also thank Dr. Karen Guillemin (University of Oregon), Dr. Raghuveer Parthasarathy (University of Oregon), and Dr. Katherine Xue (University of California, Irvine) for thoughtful discussions and constructive feedback on the manuscript.

## Notes

### Competing Interest Statement

The authors have declared no competing interest.

